# Asymptotically exact fit for linear mixed model

**DOI:** 10.1101/2023.10.25.563975

**Authors:** Yongtao Guan, Daniel Levy

## Abstract

The linear mixed model (LMM) has become a standard in genetic association studies to account for population stratification and relatedness in the samples to reduce false positives. Much recent progresses in LMM focused on approximate computations. Exact methods remained computationally demanding and without theoretical assurance. The computation is particularly challenging for multiomics studies where tens of thousands of phenotypes are tested for association with millions of genetic markers. We present IDUL and IDUL^†^ that use iterative dispersion updates to fit LMMs, where IDUL^†^ is a modified version of IDUL that guarantees likelihood increase between updates. Practically, IDUL and IDUL^†^ produced identical results, both are markedly more efficient than the state-of-the-art Newton-Raphson method, and in particular, both are highly efficient for additional phenotypes, making them ideal to study genetic determinants of multiomics phenotypes. Theoretically, the LMM likelihood is asymptotically uni-modal, and therefore the gradient ascent algorithm IDUL^†^ is an asymptotically exact method. A software package implementing IDUL and IDUL^†^ for genetic association studies is freely available at https://github.com/haplotype/IDUL.

## 1 Introduction

Genome-wide association studies (GWAS) play a pivotal role in identifying genetic variants associated with diverse traits and diseases. A key challenge in these studies is controlling for population stratification and relatedness in the sample, confounding factors, which, if unaddressed, can lead to false-positive associations. To tackle this issue, the linear mixed model (LMM) has emerged as the standard analytical approach. Earlier work focused on feasibility, as exemplified by TASSEL (Yu et al., 2006) and EMMA (Kang et al., 2008). Later works focused on improving efficiency, as exemplified by EMMAX (Kang et al., 2010), P3D (Zhang et al., 2010), FaST-LMM (Lippert et al., 2011), and GEMMA (Zhou and Stephens, 2012). More recent work, such as BOLT-LMM (Loh et al., 2015) and fastGWA (Jiang et al., 2019), aimed to make the computation feasible for large biobank datasets. One particular setting that requires high efficiency in fitting the linear mixed model is multiomics analysis, where tens of thousands of phenotypes are tested for association with millions of genetic markers.

There are two ways to improve efficiency, one is approximate computation. EMMAX and P3D fit LMM under the null and used parameters estimated from the null model for all SNPs without fitting LMM for each SNP. Svishcheva et al. (2012) approximated a SNP-specific weight with a so-called GRAMMAR-Gamma factor that is shared between SNPs, which effectively performed genomic control internally. Since the factor can be computed efficiently, this approach reduces the computation of score test from quadratic to linear (in sample sizes). BOLT-LMM framed the standard LMM in a Bayesian whole genome regression, and used a variational approximation to fit Bayesian linear regressions with Gaussian mixture priors (Loh et al., 2015). FastGWA (Jiang et al., 2019) combined three approximations: one involves fitting the LMM once under the null and using it for all SNPs; the second adopts the GRAMMAR-Gamma approach in computing score test statistics; and the third uses hard thresholding to make the kinship matrix sparse, which allows fast evaluation of the likelihood function that paves the way for a grid-search method to fit the LMM. All these approximations produced different ranking of test statistics compared to the exact computation and hence a potential power loss (more details below).

Another way to improve efficiency is through algorithmic innovation while maintaining exact computation. (Here *exact* means without aforementioned approximations; it is also short for asymptotically exact, to be discussed below.) Exact computation removes the need to consider which approximate computation works best for a given dataset (Zhou and Stephens, 2012). As an exact method, FaST-LMM first rotates the genotypes and phenotypes according to eigenvectors of the genetic relatedness matrix so that the rotated data become uncorrelated, and then optimizes a single parameter in the variance component using Brent’s method. The rotation reduces computation complexity in optimization. Another exact method GEMMA also rotates genotypes and phenotypes during optimization, but it does so implicitly. The innovation of GEMMA is its ability to evaluate the second derivatives, so that the Newton-Raphson method can be used for optimization, which converges faster than Brent’s method. The comparison between FaST-LMM and GEMMA can be found in Zhou and Stephens (2012). The Newton-Raphson method suffers from inconsistency when the initial values are distant from the optimum, to overcome this inconsistency, GEMMA starts its optimization iterations with Brent’s method and then switches to the Newton-Raphson method.

In this paper, we present IDUL and IDUL^†^ that use an iterative dispersion update to fit LMMs in genetic association studies, where IDUL^†^ is designed to be a gradient ascent algorithm by insisting on a likelihood increase between IDUL updates. We demonstrate that IDUL and IDUL^†^ are consistent and much more efficient than the state-of-the-art Newton-Raphson method in fitting LMMs, and that both are highly efficient for additional phenotypes, and thus well suited to study genetic determinants of multiomics phenotypes. Most importantly, we show that the LMM likelihood is asymptotically unimodal, and consequently IDUL^†^, a gradient ascent algorithm by design, is asymptotically exact.

## 2 Results

### 2.1 The model and the rotation

Consider a standard linear mixed model

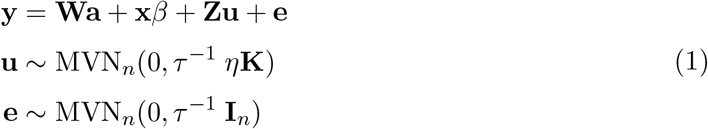

where **W** contains conventional covariates such as age and sex, including a column of 1, **x** contains genetic variant(s) to be tested for association, **u** is the random effect with **Z** as its loading matrix and kinship **K** as its covariance (both **Z** and **K** are known), MVN_*n*_ denotes an n-dimensional multivariate normal distribution, **I**_*n*_ is n-dimensional identity matrix. Denote **X** = (**W, x**) and **b** = (**a**, *β*), then **Xb** is the fixed effect, and we assume **X** has a full rank *c*. In genetic association studies, the random effect **Zu** is a nuisance term that absorbs part of the phenotype **y** that is attributable to population stratification and relatedness. We aim to find the maximum likelihood estimate (MLE) of *η*, which is the ratio between two dispersion terms (random effect **u** and random noise **e**), and conditioning on 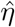 we can test the null hypothesis *β* = 0.

Denote **G** = **ZKZ**^*t*^ and its eigen decomposition **QDQ**^*t*^ (such that **QQ**^*t*^ = **I**_*n*_) where *j*-th column of **Q** is an eigenvector whose corresponding eigenvalue is the *j*-th diagonal element of the diagonal matrix **D**. Rotate both sides of (1) by multiplying **Q**^*t*^ to get

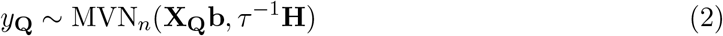

where **X**_**Q**_ = **Q**^*t*^**X, y**_**Q**_ = **Q**^*t*^**y**, and **H** = *η***D** + **I**_*n*_ is a diagonal matrix.

### 2.2 The IDUL algorithm

The iterative dispersion update for linear mixed model (IDUL) algorithm follows: S0 Initialize *η* and specify a desired precision threshold *ϵ*.

S1 For a given *η* compute **H** = *η***D** + **I**_*n*_, fit (2) using weighted least squares with weight

**H**^−1^ to obtain residual **r**, and compute 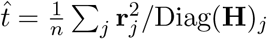.

S2 Fit **r**^2^ ∼ MVN_*n*_(*µ* + Diag(**D**)*γ, τ*^−1^**H**^2^) using weighted least squares with weight **H**^−2^ to obtain 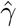 and 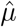, and compute 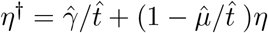.

S3 If |*η*^†^ − *η*| *< ϵ*, goto S4. Otherwise, update *η* ← *η*^†^, and goto S1.

S4 Finish and output *η*.

Intuitively, the update is mostly informed by the different level of dispersion of the residual **r**, and hence the name of the algorithm. IDUL is easy to implement; both rotation and iterative updates require only several lines of code in R (Appendices).

### 2.3 Analytic update of the IDUL

IDUL is equivalent to the following analytical update:

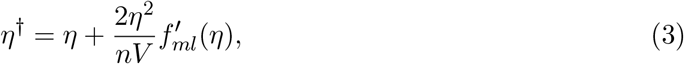

where *V* = tr(**H**^−2^)*/n* − tr(**H**^−1^)^2^*/n*^2^ is a function of *η* and 0 *< V <* 1, and 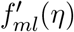 is the first derivative of the log-likelihood and can be computed analytically (Data and Methods). Note Equation (3) is derived here to study the analytic properties of the IDUL algorithm, not meant to replace step S2 in the algorithm. We make the following observations on update (3). First, when 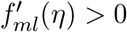 IDUL increases *η*, and when 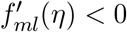 IDUL decreases *η*, until it converges to 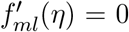, which is a local optimum. Second, Taylor expansion of the log-likelihood at *η* to get

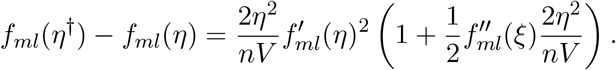

Although the factor 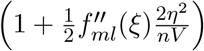 is likely to be positive (Data and Methods), there is no guarantee.

### 2.4 The IDUL^†^ algorithm

We therefore modify the algorithm and only update *η* ← *η*^†^ when the likelihood increases, and if likelihood decreases, we successively halve the step size until the likelihood increases. This technique, successive over-relaxation, is often used in iterative methods (c.f. Zhou and Guan, 2019).

R0 Initialize *η* and specify a desired precision threshold *ϵ*.

R1 With input *η*, compute **H** = *η*^†^**D**+**I**, fit (2) using weighted least squares with weight **H**^−1^ to obtain residual **r**, compute 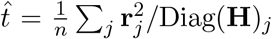 and 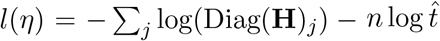.

R2 Fit **r**^2^ ∼ MVN_*n*_(*µ* + Diag(**D**)*γ, τ*^−1^**H**^2^) using weighted least squares with weight **H**^−2^ to obtain 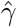 and 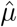, and compute 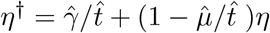.

R3 If |*η*^†^ −*η*| *< ϵ* goto R4; otherwise, do R1 with input *η*^†^ and obtain *l*(*η*^†^), and if *l*(*η*^†^) *> l*(*η*), update *η* ← *η*^†^, goto R2; otherwise update 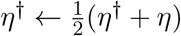, goto R3.

R4 Finish and output *η*.

The IDUL^†^ algorithm is a gradient ascent algorithm by design. Since the likelihood is bounded and the sequence of the likelihood is non-decreasing, by the standard Monotone Convergence Theorem, the IDUL^†^ algorithm must converge to a local optimum *η*^∗^ such that 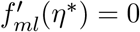. IDUL^†^ is also easy to implement in R (Appendices).

### 2.5 Asymptotically uni-modal

At an optimum such that 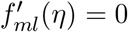, the second derivative can be simplified to the following form

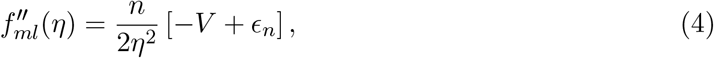

where *V* is defined in (3) and both mean and variance of *ϵ*_*n*_ varnish linearly (proportional to 1*/n*) as *n* increases (details in Data and Methods). In other words, at the local optimum, 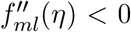 asymptotically almost sure, or with probability 1. This asymptotically local concaveness implies that with a sufficiently large sample size, the log-likelihood *f*_*ml*_(*η*) attains its unique global maximum at 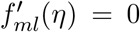. If to the contrary there are at least two local maxima, then owing to smoothness of the likelihood function Equation (5) and its derivative Equation (7), there must exist a minimum *η*^∗^ such that 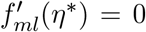 but with 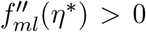, which produces a contradiction. (Intuitively, there must be a valley between two locally concave peaks, and the valley violates local concaveness.) Therefore, the likelihood function is unimodal asymptotically almost sure (or with probability 1).

### 2.6 Asymptotically exact

The notion that “an iterative method cannot be exact” (a much appreciated feedback from a reader) is false. For example, the Euclidean algorithm, used to find greatest common divisor between two integers, is an iterative algorithm, and it is exact. Another example is the Banach fixed-point-theorem, which guarantees that, when certain conditions are satisfied, fixed-point iterations always converge to a fixed point, no matter where the iteration starts. A convergence sequence is precise to an arbitrary precision and thus exact. The IDUL^†^ is a gradient ascent algorithm, and if the likelihood is unimodal, then IDUL^†^ updates produce an convergence sequence, and therefore IDUL^†^ is exact. Since the likelihood is asymptotically almost sure unimodal, so IDUL^†^ is asymptotically exact.

### 2.7 Connection with Newton-Raphson

With a sufficiently large number of samples and *η* near the optimum, *ϵ*_*n*_ in Equation (4) can be safely ignored. The analytic update of the IDUL then becomes 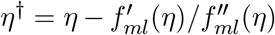, which is the Newton-Raphson method. And the IDUL^†^ becomes Newton-Raphson with successive over-relaxation. But IDUL and IDUL^†^ require no computation of the second derivative, which are expensive, and outside the neighborhood of the optimum, where the Newton-Raphson method is known to be numerically unstable (Burden and Faires, 2010), IDUL and IDUL^†^ are stable and consistent (below).

### 2.8 Consistency of IDUL

Since IDUL^†^ is asymptotically exact, it is consistent over different starting points. We numerically study IDUL’s consistency and compare it with that of the Newton-Raphson method. The genotype and phenotype datasets we used for comparison are described in Data and Methods. For each phenotype, we fitted LMM using IDUL and Newton-Raphson with the same sets of initial values. To generate initial values, we chose four non-overlapping segments from the unit interval, namely, *V*_1_ = (0.01, 0.25), *V*_2_ = (0.25, 0.5), *V*_3_ = (0.5, 0.75), and *V*_4_ = (0.75, 0.99), and for each phenotype we drew *h*_0_ uniformly from each segments to produce initial value *η*_0_ = *h*_0_*/*(1 − *h*_0_).

It takes IDUL on average 7.3 iterations to converge for each combination of phenotype and initial value, compared to 12.5 for Newton-Raphson. We compare the consistency of fitted 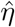 when initial values *η*_0_ were drawn from different *V* segments. IDUL had perfect consistency among four sets of initial values (Figure 1 lower triangle); Newton-Raphson did not (Figure 1 upper triangle). Taking IDUL estimates as the truth, Newton-Raphson made one error when initial values were generated from *V*_1_, 19 errors from *V*_2_, 58 errors from *V*_3_, and 70 errors from *V*_4_ (Figure 1 Diagonal). The erroneous estimates were either 0 or 1 and the proportion of 1 increases as the initial values increase (Figure 1 Diagonal). These observations are consistent with Newton-Raphson’s dependence on good initial values and that a long near flat likelihood tend to fail Newton-Raphson. Similar patterns of inconsistency of Newton-Raphson were also observed with phenotypes simulated from diverse populations in the 1000 Genomes project (Supplementary Figure S1). To overcome the inconsistency, implementing the Newton-Raphson method to fit LMM requires multiple runs from different starting points. As a comparison, IDUL and IDUL^†^ only need to run once for each model fitting, each run takes fewer iterations, and each iteration requires less computation.

**Figure 1:**
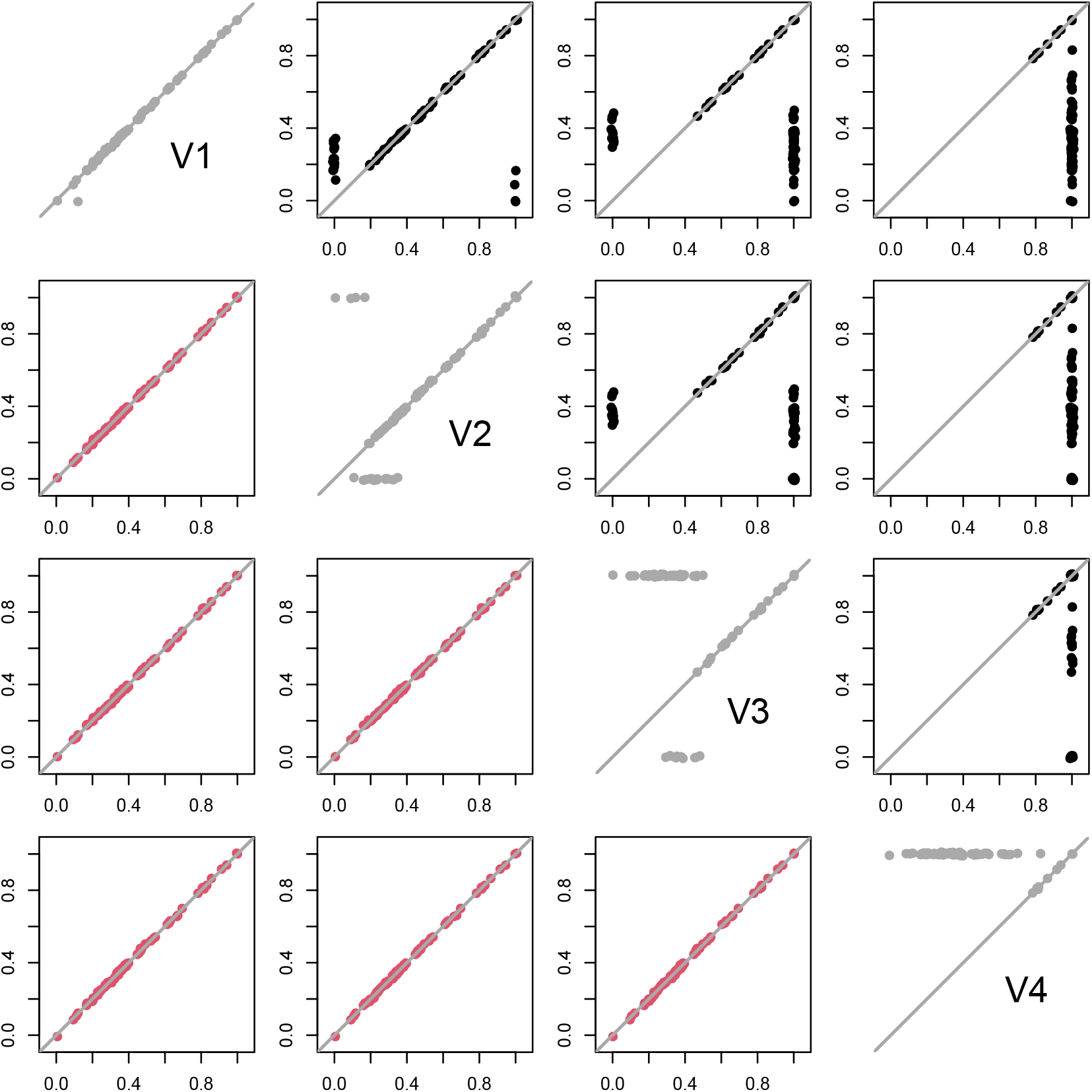
Consistency of IDUL and Newton-Raphson w.r.t different initial values. IDUL and Newton-Raphson were run on the same sets of initial values. For each phenotype, four initial values were generated with seeds randomly selected from four segments. So both IDUL and Newton-Raphson produced 4 columns of estimates each with 79 rows. The pairwise plots of the four columns of IDUL estimates were in the lower-triangle and colored in red. Those of Newton-Raphson estimates were in the upper-triangle and in black. The four diagonal plots (in gray) showed consistency or lack of it between IDUL and Newton-Raphson for different set of initial values. Estimates 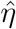 were transformed to 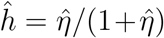 for plotting so that different panels are on the same scale. Points are jittered slightly by adding random noises for clarity.

### 2.9 Efficiency of IDUL and IDUL^†^

We implemented IDUL and IDUL^†^ into a software package IDUL that fits the LMMs and computes test statistics, and compared with results from GEMMA, a software package that fits LMMs using the Newton-Raphson method primed by the Brent’s method. The datasets we used for comparison were described in Data and Methods. We compared the likelihood ratio test (LRT) and the Wald test p-values between different methods, where the LRT requires maximum likelihood estimates of *η* and the Wald test requires REML estimates of *η* (Data and Methods). For both the LRT and the Wald test p-values, IDUL, IDUL^†^, and GEMMA reached almost perfect agreement (Supplementary Figures S2 and S3). Since IDUL^†^ is asymptotically exact, these results suggested that both IDUL and GEMMA are practically exact methods.

Both IDUL and IDUL^†^ are much more efficient than GEMMA (Table 1). For a single phenotype, GEMMA with maximum 112 threads used about 76.5 minutes for the LRT, compared with 13.6 to 15.3 minutes of IDUL and IDUL^†^ with either 32 and 64 threads. So for LRT, IDUL and IDUL^†^ are at least five times as efficient as GEMMA. GEMMA with maximum 112 threads used 98.9 minutes for Wald test, compared with 13.4 to 15.7 minutes for IDUL and IDUL^†^ with either 32 or 64 threads. So for Wald test, IDUL and IDUL^†^ is six or seven times as efficient as GEMMA.

**Table 1:** Times (minutes) used for IDUL/IDUL^†^ (version 0.81) and GEMMA (version 0.98.5) to process 1, 10, and 79 phenotypes. GEMMA used maximum 112 threads and IDUL/IDUL^†^ used 64 and 32. LRT: likelihood ratio test, which requires maximum likelihood estimates of *η*. Wald: Wald test, which requires REML estimates of *η*. Taking multiple phenotypes to analyze one by one was not implemented in GEMMA, and its wall time for 10 and 79 phenotypes are missing.

| LRT |  |  | Methods |  | Wald |  |  |
| --- | --- | --- | --- | --- | --- | --- | --- |
| $p = 1$ | 10 | 79 | Algo | Threads | $p = 1$ | 10 | 79 |
| 76.5 | - | - | GEMMA | 112 | 98.9 | - | - |
| 13.6 | 14.9 | 19.3 | IDUL | 64 | 13.4 | 16.8 | 24.5 |
| 13.9 | 14.7 | 19.2 | IDUL<sup>†</sup> | 64 | 13.9 | 16.5 | 24.6 |
| 15.3 | 16.4 | 23.8 | IDUL | 32 | 15.7 | 18.8 | 31.9 |
| 14.9 | 15.9 | 25.2 | IDUL<sup>†</sup> | 32 | 15.1 | 19.7 | 33.0 |

Most remarkably, IDUL and IDUL^†^ are highly efficient for additional phenotypes. Taking LRT and 64 threads as an example, IDUL and IDUL^†^ only spent about one extra minute (about 8% of time spent for the first phenotype) to compute like ratio test for additional 9 phenotypes, and less than six extra minutes (about 40% of time spent for the first phenotype) for additional 78 phenotypes. Similar high efficiency for additional phenotypes was also observed with the Wald test and with 32 threads (Table 1). Also note that doubling the number of threads used by IDUL/IDUL^†^ resulted in a small improvement in speed for a single phenotype, but a larger improvement with additional phenotypes. The high efficiency for extra phenotypes makes IDUL/IDUL^†^ ideal to study genetic determinants of multiomics phenotypes.

### 2.10 Exact vs approximation

The IDUL^†^ is an asymptotically exact method. Zhou and Stephens (2012) classified methods into approximate methods and (practically) exact methods and demonstrated that 1) among exact methods available at the time, GEMMA is most efficient, outperforming FaST-LMM, which in turn outperforms EMMA by an order of magnitude; and 2) approximate method such as EMMAX, which uses the parameter estimated under the null to compute test statistics for all SNPs, evidently biased test statistics in some dataset. A recent approximate method, fastGWA by (Jiang et al., 2019), in addition to other approximations, applied hard thresholding on kinship matrix *K* in Equation (1) to make it sparse and exploited the sparsity in fitting the linear mixed model and computing test statistics. The approximation makes fastGWA capable of analyzing large datasets such as UK Biobank. For multiomics datasets such as that of the Framingham Heart Study, however, the hard thresholding approach appears less satisfactory, presumably due to closer relatedness between samples and larger effect sizes in multiomics dataset. Figure 2 shows a comparison of test statistics of four protein phenotypes between exact computation and hard thresholding approximation. The inconsistency can be rather pronounced for some phenotypes, suggesting potential difficulty with approximate methods in multiomics data for closely related samples such as in the Framingham Heart Study.

**Figure 2:**
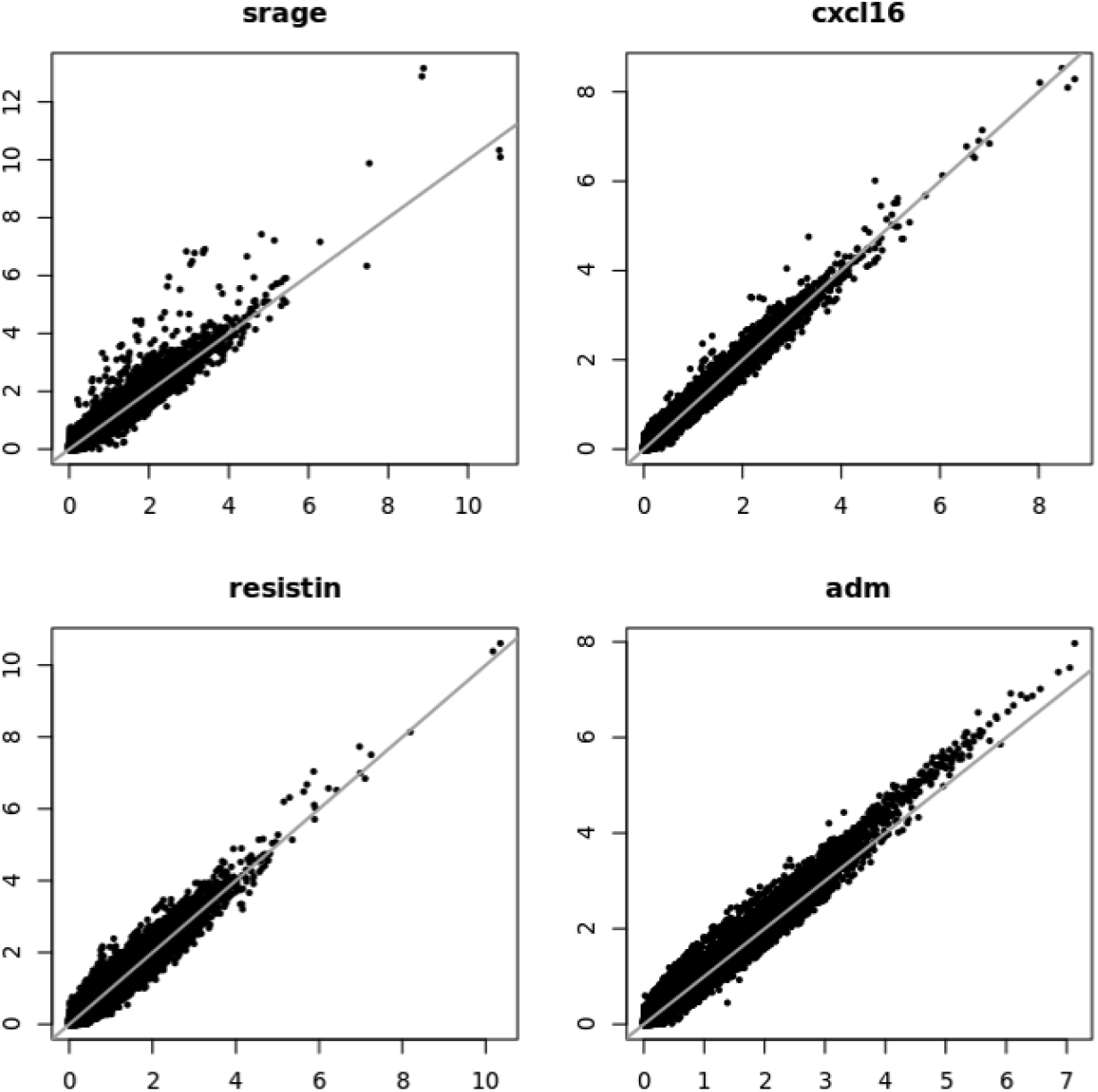
Comparison between exact and approximate method. The test statistics are −log_10_ p-values, with the exact statistics on x-axis, and the approximate statistics on y-axis. The genomic control values for the exact methods are 0.998, 0.997, 1.013, 1.045 vs 1.036, 1.007, 1.043, 1.066 for the approximate method.

## 3 Discussion

In this paper we documented two novel methods IDUL and IDUL^†^ to fit linear mixed model in the context of genetic association studies. IDUL^†^, a modification of IDUL, is a gradient ascent algorithm by design. Both IDUL and IDUL^†^ are much more efficiency than the state-of-the-art Newton-Raphson method. The fundamental contribution of the paper, however, is that the log-likelihood of the linear mixed model is asymptotically locally concave at the optimum (we hypothesized that the same is true for REML likelihood, see Data and Methods). Consequently, the likelihood of the standard linear mixed model is asymptotically uni-modal. Therefore, IDUL^†^, a gradient ascent algorithm, is an asymptotically exact method.

We demonstrated that IDUL and IDUL^†^ are much more efficient than the Newton-Raphson method in fitting the standard LMM. We would like to point out that IDUL and IDUL^†^ are specialized algorithms that take advantage of a specific dispersion structure only available in limited settings such as the LMMs. Newton-Raphson, on the hand, is a general method that can be applied in many settings. In addition, our theoretical analysis relies on the assumption of normality of the phenotypes. Although the assumption can perhaps be relaxed to having finite first and second moments, such as binary phenotypes, it is prudent to examine phenotypes and perform quantile normalization when they show severe departure from normality.

When there are population structures among the samples, such as in the simulation studies using 1000 Genomes datasets, IDUL and IDUL^†^ updates oscillate like a damping pendulum. The algorithms still converge, but the oscillation increases the number of iterations from several to several dozen. This oscillation can be resolved by controlling for leading eigenvectors (such as top three PCs). It can also be resolved by making IDUL and IDUL^†^ lazy. Specifically, the update 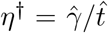 has a step size that is a fraction (specifically tr(**H**^−1^)*/n*) of the step size of that IDUL update 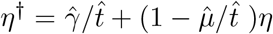. We can take an average of the two updates to get 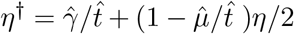 (more details in Appendices), and this lazy update brings oscillation to a quick stop (Supplementary Figure S4).

IDUL is designed with genetic association with multiomics data in mind, where the same set of genotypes are tested against thousands or tens of thousands phenotypes for association, where rotation of genotype vectors (left multiplying *Q*^*t*^) only needs to be done once for all phenotypes. Suppose in a study we have *n* sample, *m* SNPs, *p* phenotypes, and *c* covariates, then the total complexity is *O*(*n*^3^ + (*m* + *p* + *c*)*n*^2^ + *tmp*(*nc*^2^ + *c*^3^)), where *O*(*n*^3^) is for eigen decomposition, *O*((*m* + *p* + *c*)*n*^2^) is for rotation, and *O*(*tmp*(*nc*^2^ + *c*^3^)) is for model fitting of all SNP-phenotype pairs, where *O*(*nc*^2^ + *c*^3^) is the complexity of linear regression, and we assume IDUL converges on average in *t* iterations. For a typical study where *m > p > n* ≫ *c*, the dominate term in total complexity is *O*(*tmpnc*^2^), which is linear in the size of the study, namely, *n, m*, and *p*. IDUL^†^ has the same complexity as IDUL, with a slightly larger equivalent *t* for extra computation to evaluate likelihood. The Newton-Raphson method also has the same complexity, but it tends to have a much larger equivalent *t* than IDUL and IDUL^†^ because essentially it needs to run multiple times to compensate for its lack of consistency, each run takes more iterations to converge, and each iteration takes more computation because it requires second derivatives. Table 1 in fact confirmed this intuition.

This strategy of reusing intermediate computation of each genetic variant for multiple phenotypes was also a feature in Regenie (Mbatchou et al., 2021), a whole genome regression (WGR) approach to the linear mixed model that is comparable to BOLT-LMM. Compare to WGR, the standard linear mixed model is more flexible in its applications. For example, in testing for parental origin effect and/or controlling for local ancestry, the standard model only needs to add extra variates and covariates, while the rest of the computation remains the same. But WGR has to change model and priors, and perhaps the details on computation, to make it happen. Strictly speaking, WGR is not a standard linear mixed model. For example, the standard linear mixed model can incorporate different estimates of genetic relatedness matrix such as the one estimated by Kindred (Guan and Levy, 2024), while WGR is stuck with sample correlation as its equivalent genetic relatedness matrix.

In our application, the random effect was treated as a nuisance parameter and our goal is testing fixed effect. Under this context, the MLE is preferred over the REML estimate, because ML estimate of the fixed effects are unbiased (West et al., 2014). REML estimates, however, is preferred when the interest is the variance component, such as in estimating trait heritability, because it produces unbiased estimate of the variance component (i.e, *η*). IDUL can be adapted to obtain REML estimate based on its analytical update (Data and Methods). The standard software package to estimate heritability is GCTA (Yang et al., 2011), which employs the Average Information REML for model fitting (Gilmour et al., 1995). The software has trouble dealing with modestly small sample sizes. Using an Australian height dataset (Yang et al., 2010), we performed down-sampling at 90%, 70%, and 50% of total 3925 samples 100 times each, obtain REML estimates using IDUL for different estimates of kinship matrices without any issue (Supplementary Figure S5). As a comparison, the Average Information REML implemented in GCTA had difficulty even with 90% down-sampling and produced untenable results.

## 4 Data and Methods

### 4.1 Data sets

We used datasets from the Framingham Heart Study (FHS) to conduct numerical comparisons between IDUL and the Newton-Raphson method. Funded by the National Heart, Lung, and Blood Institute (NHLBI), the FHS includes many independent three generational pedigrees, nuclear families, trios, duos, and singletons (Kannel et al., 1979). We used 5757 samples with whole genome sequencing data through NHLBI’s TOPMed program (Taliun et al., 2021) and who also have protein immunoassays obtained through the NHLBI’s Systems Approach to Biomarker Research in Cardiovascular Disease (SABRe CVD) Initiative (Ho et al., 2018). With 79 phenotypes, this dataset represents a mini example of multiomics data.

We also used genotype data from the 1000 Genomes project (Auton et al., 2015) with simulated phenotypes to demonstrate the effectiveness of our method for diverse populations. Finally, Australia height data from Queensland Institute of Medical Research was used for down-sampling study to demonstrate the robustness of IDUL with small sample sizes.

### 4.2 Likelihood and the derivatives

Following notations in Equations (1) and (2), define projections 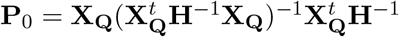 and **P** = **I**_*n*_ − **P**_0_. Denote **P**_*x*_ = **H**^−1^**P**. The marginal log-likelihood function for *η* for model 2 is

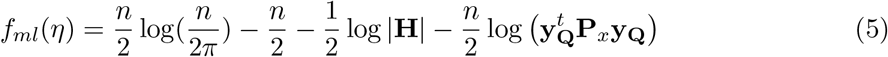

Because **H** is diagonal and Equation 17, *f*_*ml*_(*η*) can be evaluated efficiently. For log-restricted likelihood is

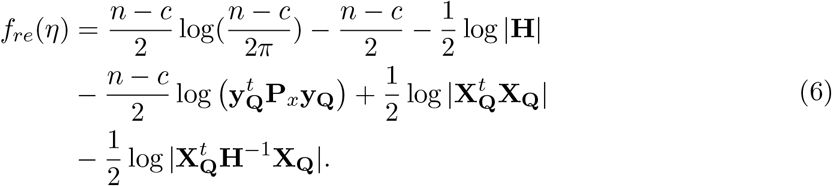

The first and second derivatives of the log-likelihood function are

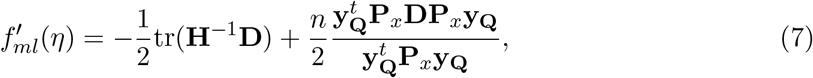

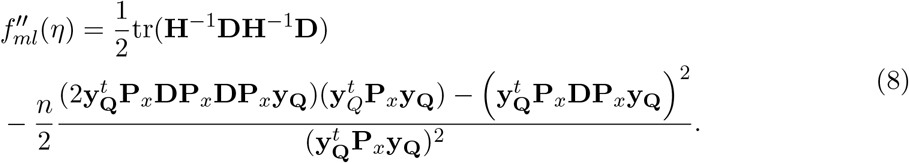

Since we work with the rotated system (2), the only matrix calculus identity needed to derive these is 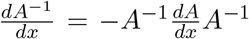 where matrix *A* is a function of a scalar *x*. For log-restricted likelihood we have

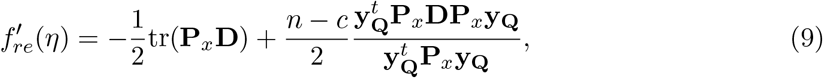

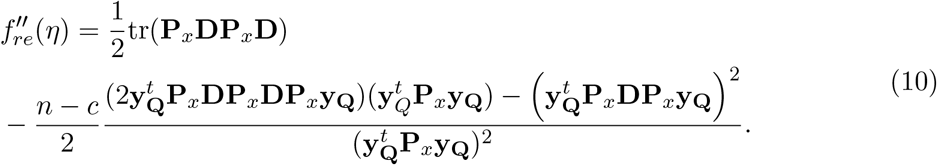

These likelihood and derivatives are in the same form as those in (Zhou and Stephens, 2012).

### 4.3 Evaluation of the first derivatives

We simplify the directive of likelihood functions using residuals from S1 of the IDUL.

#### Proposition 1.

*Let* **r** = **Py**_**Q**_, *the following hold:*

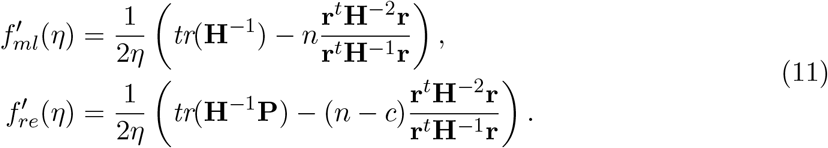

Proof of Proposition 1 is deferred to Appdendices. Because **H** is diagonal, evaluation of 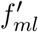 has a complexity of *O*(*n*). To evaluate 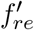 note tr(**H**^−1^**P**) = tr(**H**) − tr(**HP**_0_), while 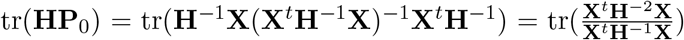. Since **X**^*t*^**H**^−1^**X** is *c × c*, its inverse has complexity *O*(*c*^3^). Therefore the complexity of evaluating 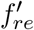 is *O*(*n* + *nc* + *c*^3^).

### 4.4 IDUL update in an analytical form

To study the theoretical property of IDUL, we derive its analytic form by computing 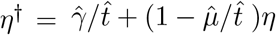 in S2 of the IDUL.

#### Lemma 2.

*The update of IDUL for maximum likelihood estimate is*

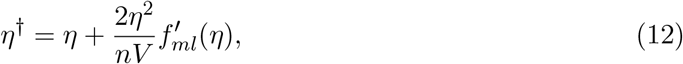

*where V* = *tr*(**H**^−2^)*/n* − *tr*(**H**^−1^)^2^*/n*^2^ *>* 0 *and is bounded, and* 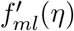 *is the derivative of the log-likelihood evaluated at η*.

Proof of Lemma 2 is deferred to Appendices. The Taylor expansion of *f*_*ml*_ at *η* is 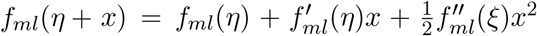 for some *ξ* ∈ (*η* − *x, η* + *x*). Let *x* = *η*^†^ − *η*, we get 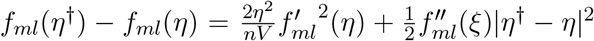. Substitute 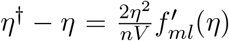 back in, we have

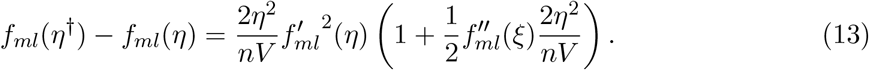

When *η* is near the optimal, we have *f*_*ml*_(*η*^†^) −*f*_*ml*_(*η*) *>* 0 for large *n* by virtue of Equation 15. But when *η* is not near optimal, there is no guarantee that the likelihood always increase. Thus we modify IDUL by evaluating and comparing likelihood to guarantee the likelihood increase between updates.

### 4.5 From MLE to REML

With maximum likelihood update (12) at hand, we can obtain REML estimate by substituting S2 in IDUL and R2 in IDUL^†^ with

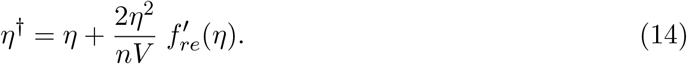

Of course, for IDUL^†^ the likelihood in R1 needs to be revised to REML likelihood.

### 4.6 Asymptotically locally concave at optimum

Finally, we quantify the second derivative of the log-likelihood function to show it is asymptotically locally concave at the local optimum.

#### Theorem 3.

*Let η be an optimum of log-likelihood function such that* 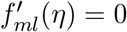, *then*

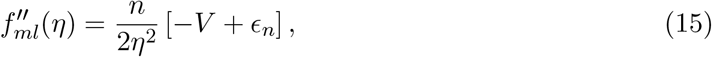

*where V* = *tr*(**H**^−2^)*/n* − *tr*(**H**^−1^)^2^*/n*^2^ *>* 0 *and is bounded, and both mean and variance of ϵ*_*n*_ *decreases linearly* 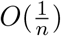. *Thus, at a local optimum such that* 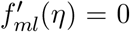, *we* 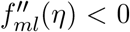 *asymptotically almost sure*.

Proofs of Theorem 3 is deferred to Appendices. Owing to the similarity of expression between 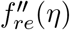 and 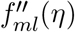, we believe the following conjecture can be proved by exploring the connections between eigenvalues of **H**^−1^**P** and **H**^−1^, evidenced by that **H**^−1^**PPv** = *λ***Pv** implies **H**^−1^**Pv** = *λ***Pv**.

#### Corollary 4.

*Let η be an optimum of REML-likelihood function such that* 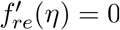, *then*

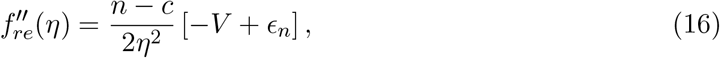

*where V* = *tr*(**H**^−1^**PH**^−1^**P**)*/*(*n*−*c*)−*tr*(**H**^−1^**P**)^2^*/*(*n*−*c*)^2^ *>* 0, *and both mean and variance of ϵ*_*n*_ *decreases linearly* 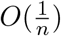. *Thus*, 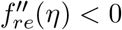*almost sure, or with probability* 1, *for a sufficiently large n*.

### 4.7 Computing association p-values

With maximum likelihood estimate and REML estimate of *η*, we can compute p-values for association. To test the null hypothesis *β* = 0, we computed the likelihood ratio test (LRT) p-values using maximum likelihood estimates as suggested by (Yu et al., 2006) and Wald test p-values using REML estimates as suggested by (Kang et al., 2008). Both test statics were described in clean detail in Supplementary of (Zhou and Stephens, 2012).

## 5 Acknowledgements

This research is supported by the Division of Intramural Research of the National Heart, Lung, and Blood Institute, Bethesda, MD (D.L. Principal Investigator). We thank Nick Martin and the Queensland Institute of Medical Research for making the Australia height data available to us.

## 6 Author contributions

Y.G. conceived the study, developed the methodology, implemented the computational software and performed experiments, analyzed results, and wrote the manuscript. D.L. supervised the work, provided access to data and computation, edited and approved the final manuscript.

## 8 Appendices

### 8.1 R code for IDUL and IDUL^†^

~~~
eG=eigen(G); #G=ZKZ^t;
D=ifelse(eG$values < 0, 0, eG$values);
xQ=t(eG$vectors) %*% cbind(W, x); #rotate covariates W and genotype x;
yQ=t(eG$vectors) %*% y; #rotate phenoytpe y;
idul=function(xQ,yQ,D,eta,epsilon) {
     repeat {
        H = eta * D + 1;
        r2 = lm(yQ∼xQ, weights=1/H)$residuals^2;
        tauinv = mean(r2/H);
        fit2 = lm(r2∼D, weights=1/H/H);
        param=fit2$coefficients;
        eta1 = max(0,param[2]/tauinv+(1-param[1]/tauinv)*eta);
        print(c(eta,eta1),digits=6);
        if(abs(eta1-eta) < epsilon) {break;}
        eta = eta1;
  };
  return(eta);
}
idul_plus=function(xQ,yQ,D,eta,epsilon) {
    H = eta * D + 1;
    r2 = lm(yQ∼xQ, weights=1/H)$residuals^2;
    tauinv = mean(r2/H);
    like = -sum(log(H)) - length(D) *log(tauinv);
    repeat {
      fit2 = lm(r2∼D, weights=1/H/H);
      param=fit2$coefficients;
      eta1 = max(0, param[2]/tauinv+(1-param[1]/tauinv)*eta);
      while(abs(eta1-eta)>epsilon){
         H1= eta1 * D + 1;
         r2 = lm(yQ∼xQ, weights=1/H1)$residuals^2;
         tauinv1 = mean(r2/H1);
         like1 = -sum(log(H1)) - length(D)*log(tauinv1);
         if(like1 >= like) {break;}
         eta1 = (eta1+eta)/2;
   }
   print(c(eta,eta1, like, like1));
   if(abs(eta1-eta) < epsilon) {break;}
   eta = eta1; H=H1; like=like1; tauinv=tauinv1;
};
 return(eta);
}
~~~

### 8.2 Proof of Proposition 1

*Proof*. We first simplify two expressions

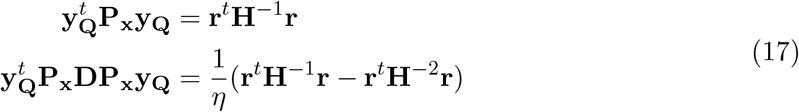

1) By **PP** = **P** and **r** = **Py**_**Q**_, we have 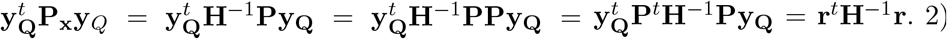 Recall **H** = *η***D** + **I**_*n*_, so 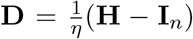, then by direct computation we have 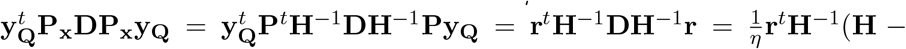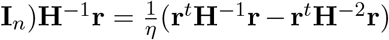. With these two reduced expressions, the first derivatives can be transformed as following:

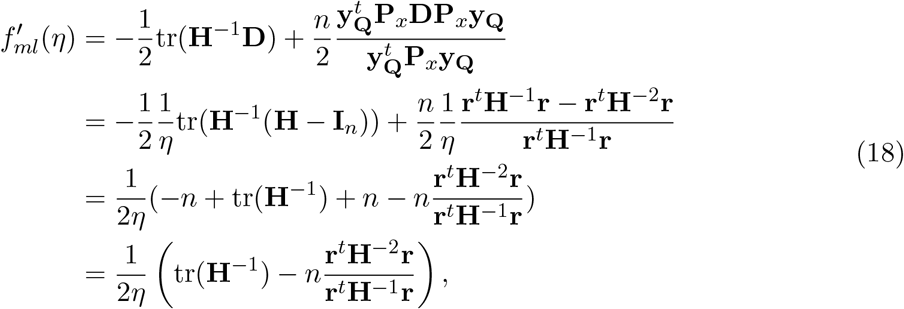

and

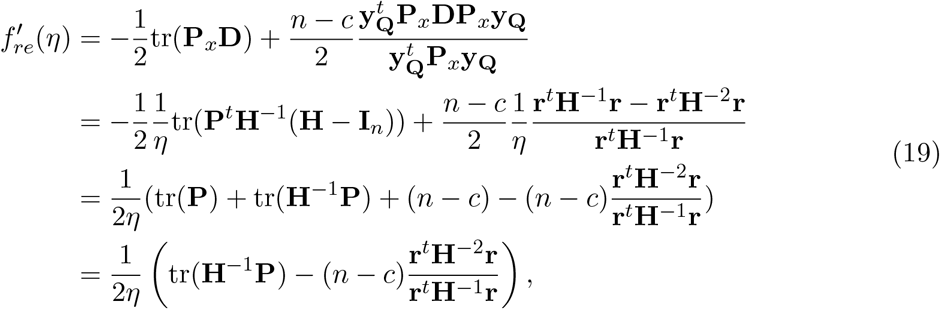

where the last equality holds because tr(**P**) = tr(**I**_*n*_) − tr(**P**_0_) and **P**_0_ is a projection with rank *c* thus tr(**P**_0_) = *c*.

### 8.3 Proof of Lemma 2

*Proof*. Step 1 is a weighed linear regression, we can compute **r** = **Py**_**Q**_. Step 2 is also a weighted linear regression with two covariates, so that its solution can be directly computed. Let vector **d** be the diagonal elements of **D** and **1** is the vector of 1 and **s** is component wise square of **r**, we have

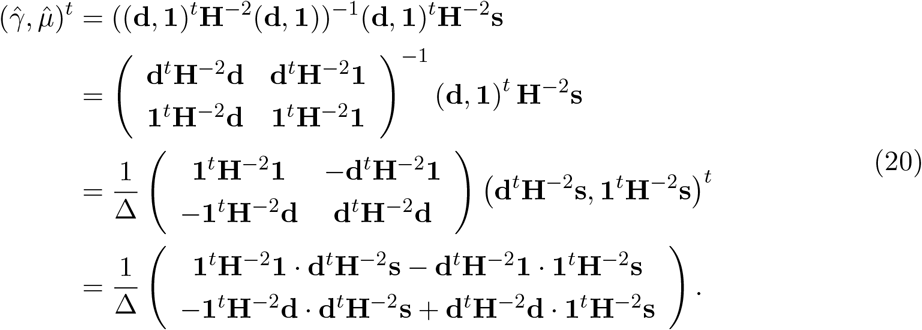

Since **H** and **D** are diagonal, we have

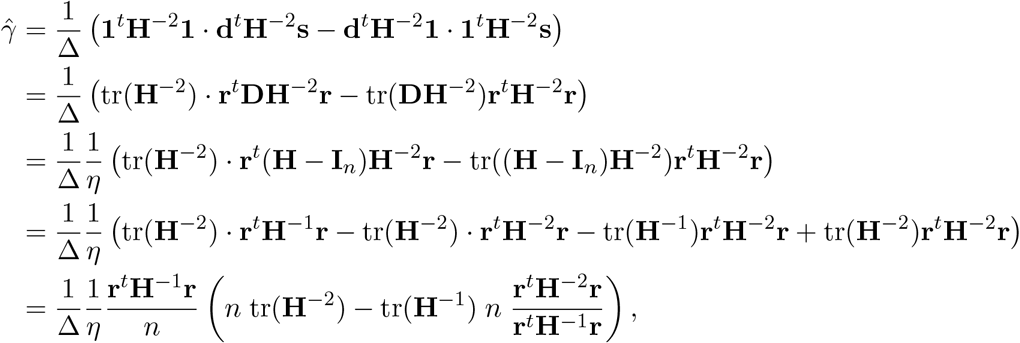

and

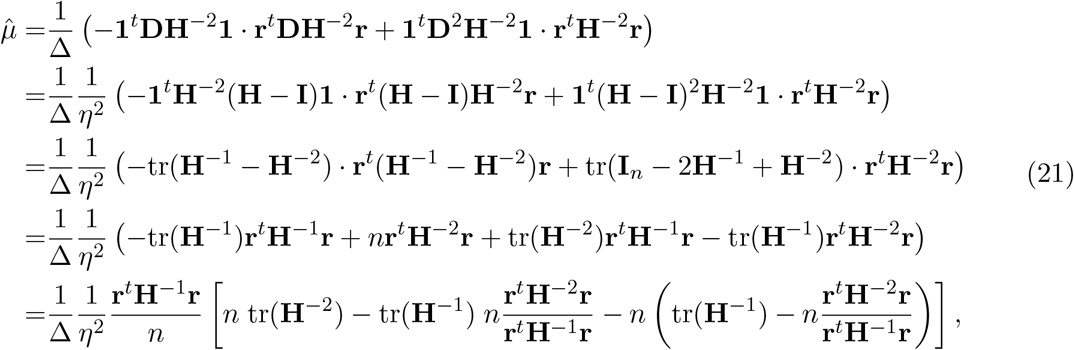

Where

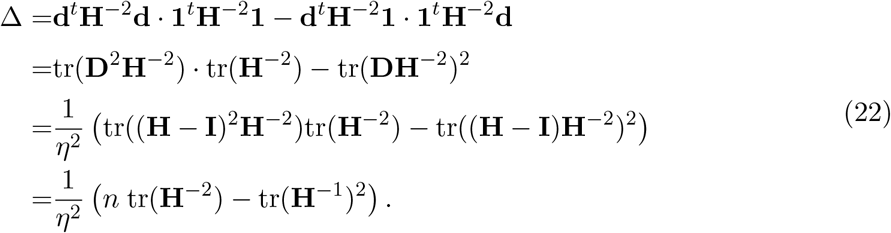

Note 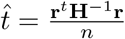 and 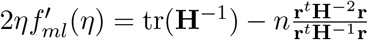, we have

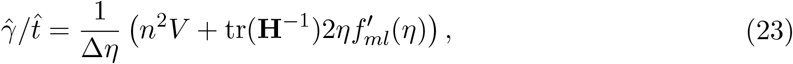

and

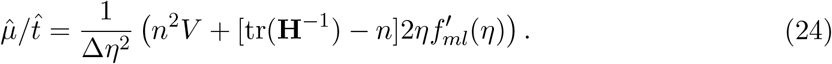

Putting together and plug in 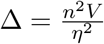 to get

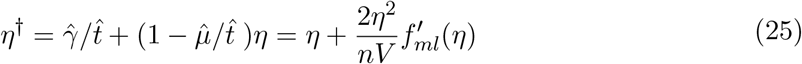

Alternatively, note

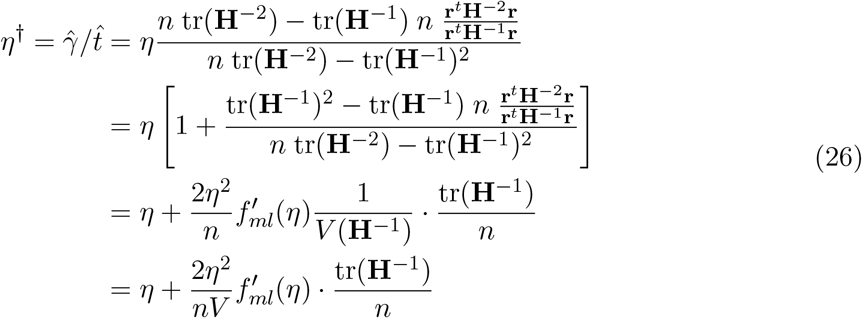

has a fractional step size with fraction being 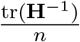, so that we can combine updates (26) and (25) to get a lazy update

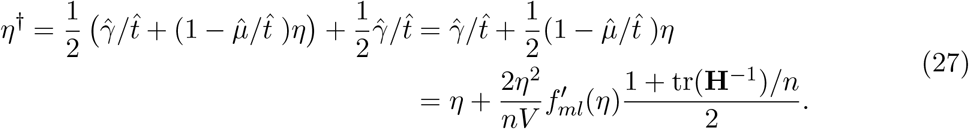

Finally, note that **H** = *η***D** + **I**_*n*_ and since *η >* 0 and **D**_*j*_ *>* 0 so **H**_*j*_ *>* 1. Denote *h*_*j*_ the *j*-th diagonal element of **H**^−1^, we have

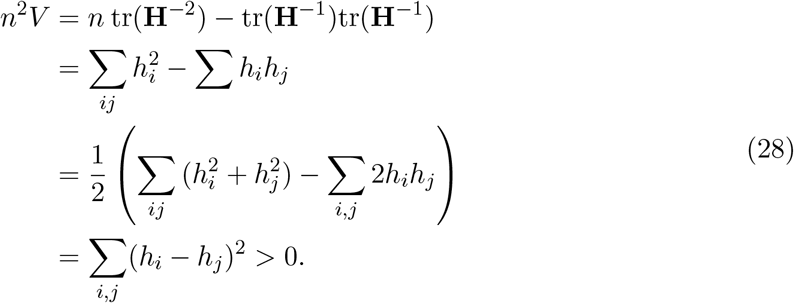

On the other hand,

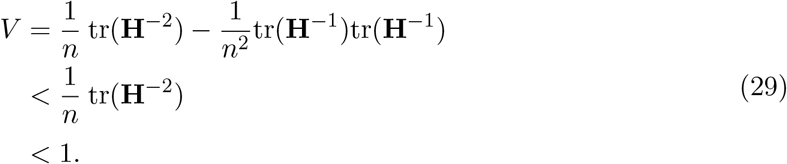

### 8.4 Proof of Theorem 3

*Proof*. We first simplify a long term 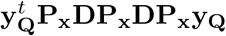. Using 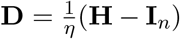 twice, we get:

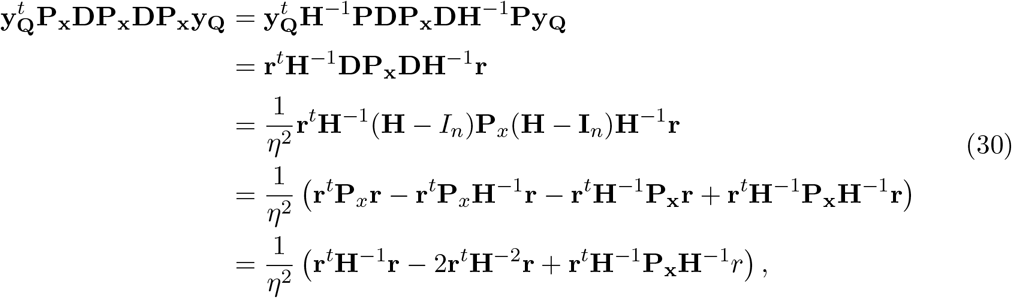

where in the last equality, we used **r**^*t*^**P**_*x*_**H**^−1^**r** = **r**^*t*^**P**^*t*^**H**^−1^**H**^−1^**r** = **r**^*t*^**H**^−2^**r**. Combing this and 17 and the assumption that 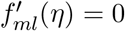 to get

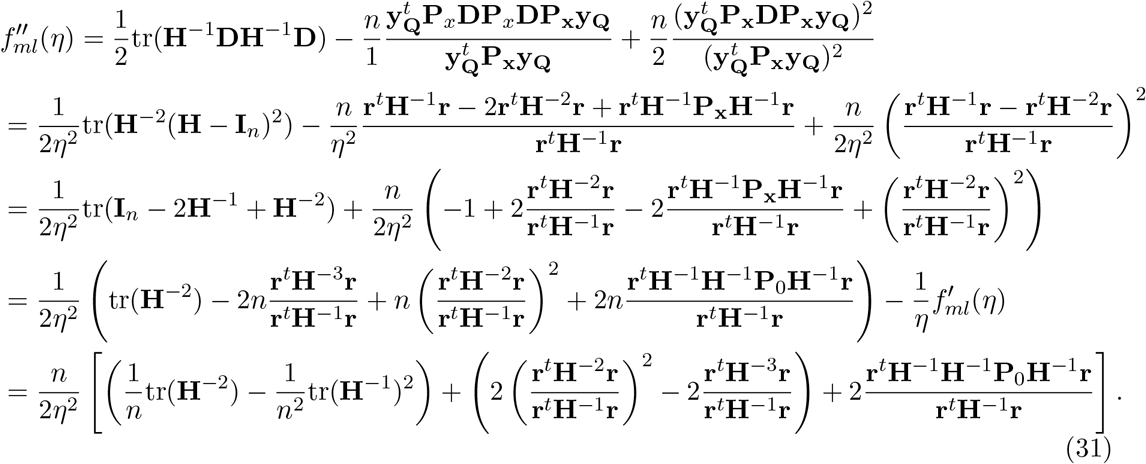

We examine three terms in the square bracket in turn. Denote *h*_*j*_ the *j*-th diagonal element of **H**^−1^, with reference to Equation 28 the first term can be transformed to

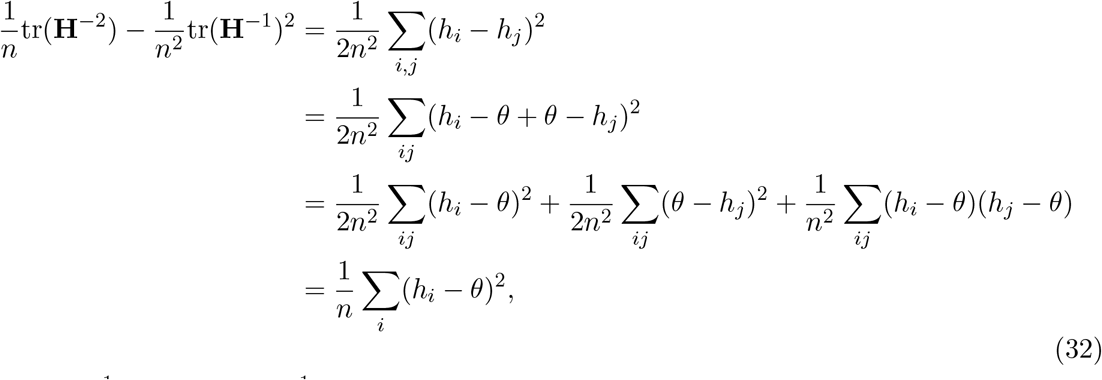

where 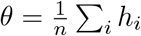 so that 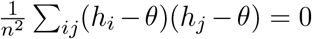. This is the average of the squared errors and we denoted it by *V* (**H**^−1^).

The second term can be transformed to

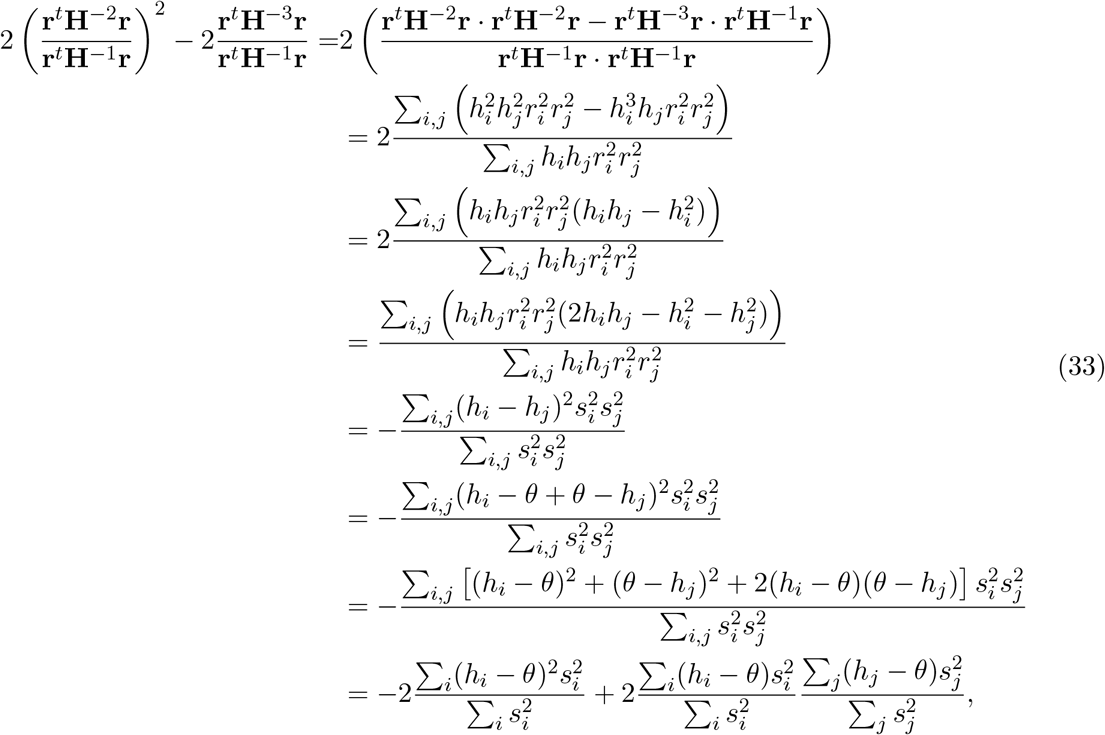

where we denote 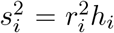 . To this end, the term 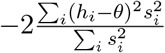 can be seen as a stochastic average of squared errors, note 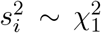, and 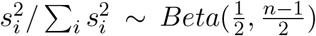 with mean 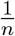 and variance 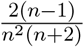 . Therefore, the mean of the term is −2*V* (**H**^−1^), and the variance of the term is 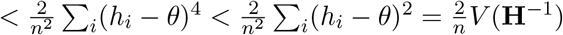, which goes to 0 as *n* increases.

Identifying the second term as a square of sum of random variables, and we apply central limit theorem to show it’s a square of a normal random variable whose mean and variance both vanish as *n* increases. Let 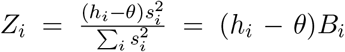, where 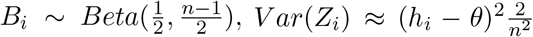. Denote *Z* = Σ_*i*_ *Z*_*i*_ and 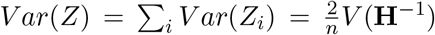. We are to apply Lyapunov central limit theorem, so let us check that Lyapunov’s condition holds:

*E*(*Z*_*i*_) = (*h*_*i*_ − *θ*)*/n* and 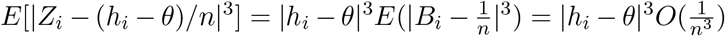, so that 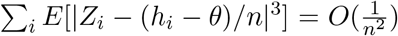, and 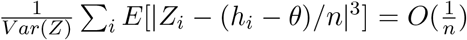, which satisfy Lyapunov’s condition. Then by Lyapunov central limit theorem 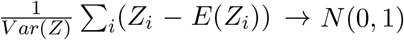, and equivalently 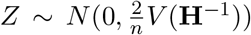. Thus, the second term has mean 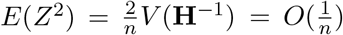, and variance (computed via scaled 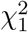) is 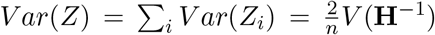, which go to 0 as *n* increases.

The third term can be transformed to

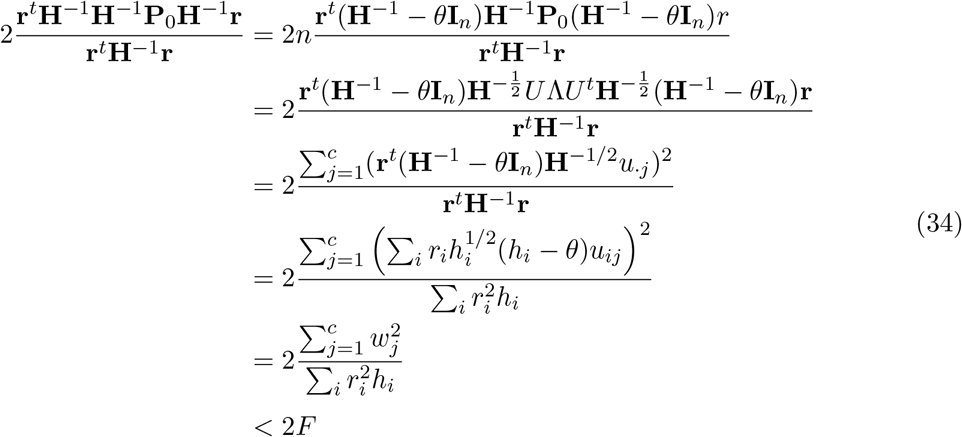

where 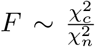 . The first equality holds because **r** = (**I**_*n*_− **P**_0_)**y**_**Q**_ and **r**^*t*^**H**^−1^**P**_0_**v** = 0 and **vH**^−1^**P**_0_**r**^*t*^ = 0 for any **v** (or in other words, we add terms equal to 0); the second equality holds because 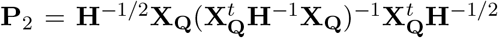 is also a projection, and **P**_2_ is symmetric and has the same rank and trace as **P**_0_, therefore **P**_2_ = *U* Λ*U* ^*t*^ where *U* is orthonormal; the third equality holds because Λ has *c* eigenvalues 1 and *n − c* eigenvalues 0; the fourth equality holds by definition; in the fifth equality, we define 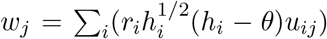, and because is standard normal, *w*_*j*_ is a weighted sum of normal random variables, and itself a normal with mean 0 and variance 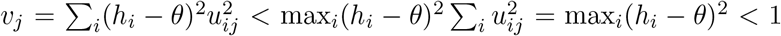, which gives the last inequality. (Note that *u*_*·j*_ is an orthonormal basis, and 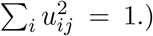 Finally, 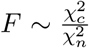, and 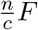 *F* follows F-distribution with d.f. *c* and *n* (*c << n*), whose mean and variance is *O*(1), and thus 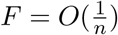 goes to 0 as *n* increases.

Putting together,

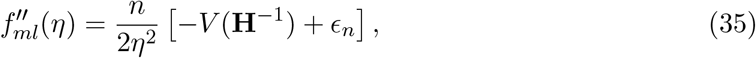

with both mean and variance of *ϵ*_*n*_ decreases linearly 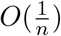. Thus, 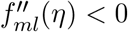 asymptotically almost sure, or with probability 1, at where 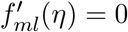.

