## Supplementary for "Asymptotically exact fit for linear mixed model"

Yongtao Guan and Daniel Levy
National Heart, Lung, and Blood Institute

March 25, 2024

### 1 Figures

#### 1.1 1000 Genomes dataset

The 1000 Genomes project contains 2504 samples from diverse populations. While Sabre dataset in the main text is an example dataset that dominated by relatedness, the 1000 Genomes samples is an example dataset that is dominated by population stratification. We simulated 100 phenotypes with nominal heritability of 0.80. For each phenotype, we fit LMM using IDUL and Newton-Raphson with the same sets of initial values. To generate initial values, we chose four non-overlapping segments from the unit interval, namely, $V_1 = (0.01, 0.25), V_2 = (0.25, 0.5), V_3 = (0.5, 0.75)$, and $V_4 = (0.75, 0.99)$, and for each phenotype we drew $h_0$ uniformly from each segment to produce initial value $\eta_0 = h_0/(1 - h_0)$.

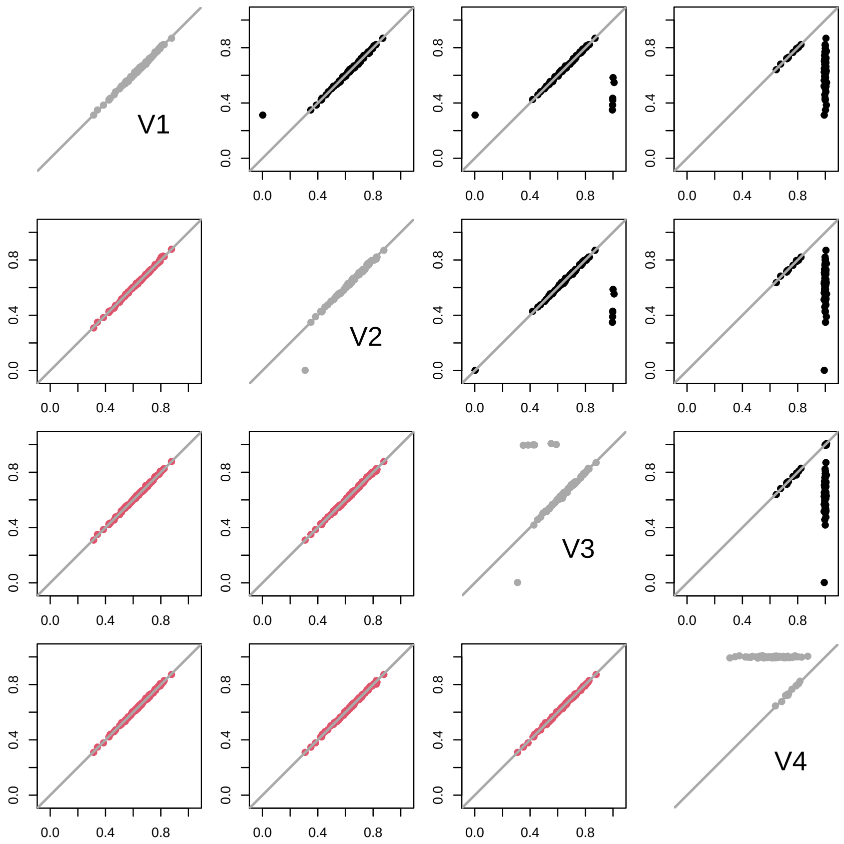


Figure S1: Consistency of IDUL and Newton-Raphson. IDUL and Newton-Raphson were run on the same sets of initial values. For each phenotype, four initial values were generated with seeds randomly selected from four segments. So both IDUL and Newton-Raphson produced 4 columns of estimates each with 100 rows. The pairwise plots of the four columns of IDUL estimates were shown in the lower-triangle and colored in red. Those of Newton-Raphson estimates were shown in the upper-triangle and in black. The four diagonal plots (in gray) showed consistency or lack of it between IDUL and Newton-Raphson for different set of initial values. Estimates $\hat{\eta}$ were transformed to $\hat{h} = \hat{\eta}/(1+\hat{\eta})$ for plotting so that different panels are on the same scale. Points are jittered slightly by adding random noises for clarity.

### 1.2 IDUL and IDUL$^\dagger$

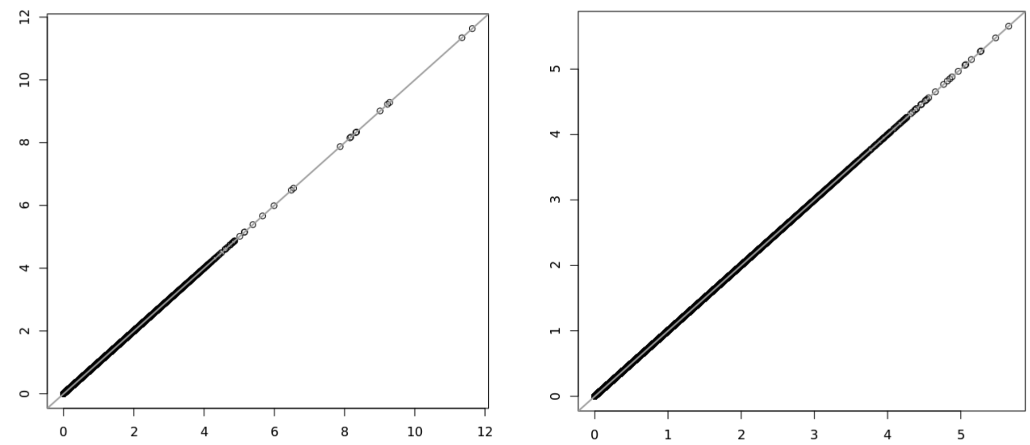


Figure S2: Comparison of $-\log_{10} p$ between IDUL (x-axis) and IDUL† (y-axis). The left panel plot Wald test p-values for a quantile normalized srage level, and the right panel is for LRT p-values for a quantile normalized apoa1 expression. IDUL and IDUL† achieved identical results.

### 1.3 IDUL and GEMMA

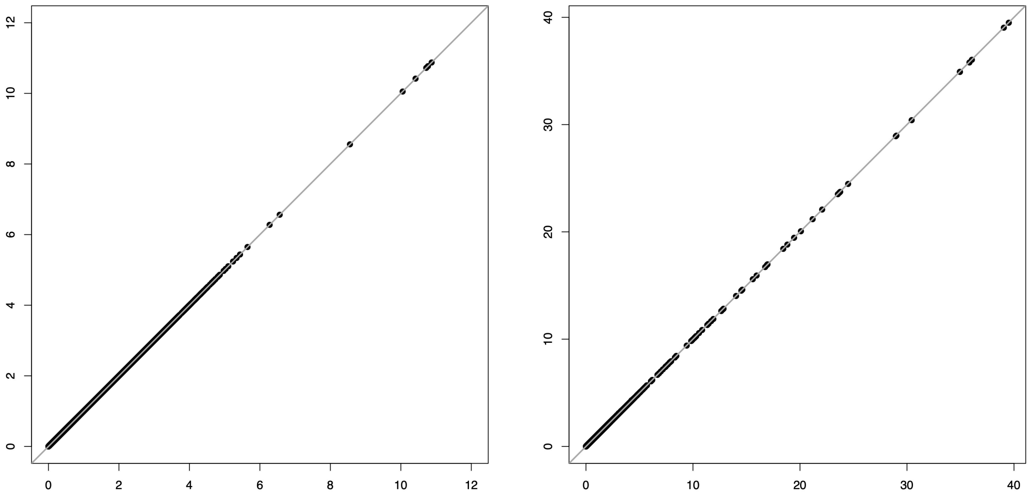


Figure S3: Comparison of $-\log_{10} p$ between IDUL (x-axis) and GEMMA (y-axis). The left panel plot Wald test p-values for a quantile normalized HDL level, and the right panel is for LRT p-values for a quantile normalized lipoprotein expression. IDUL and GEMMA achieved near identical results.

### 1.4 Lazy update

In this simulation, we used all 2504 samples from the 1000 Genomes project with simulated phenotype.

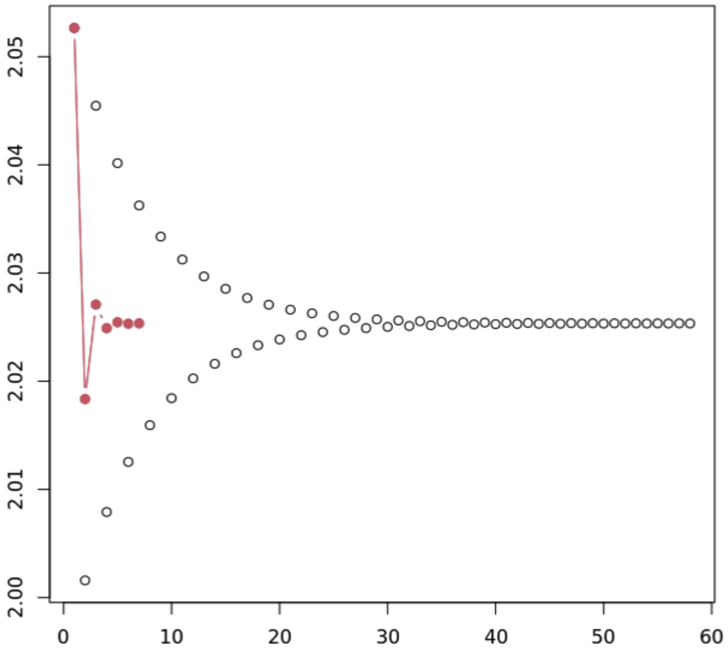


Figure S4: IDUL lazy update is effective to bring oscillation to a quick stop. The trajectory of non-lazy update was plotted in circle, and lazy update in dots.

### 1.5 IDUL is reliable for small samples.

An Australian height data was used to study stability of IDUL estimates for small sample sizes.

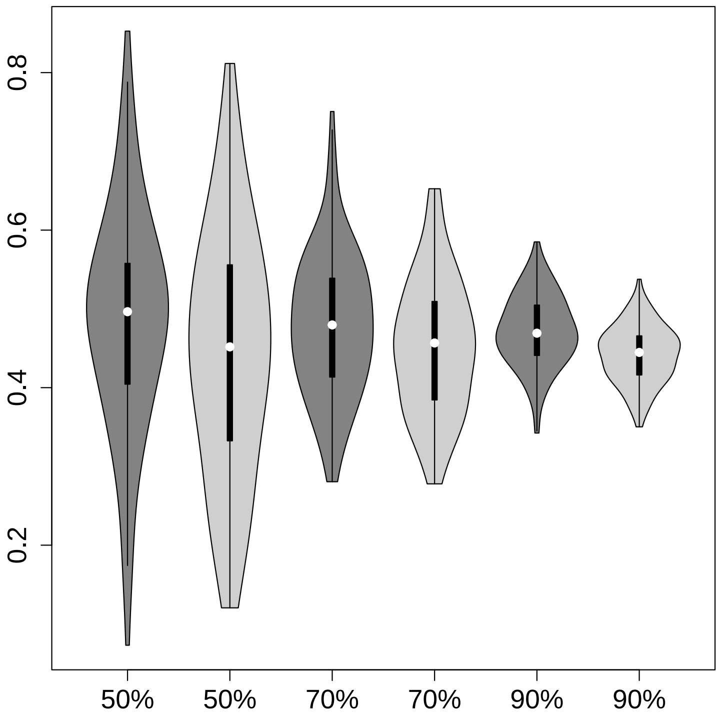


Figure S5: Violin plots of heritability estimates using IDUL. This is to show feasibility of IDUL on small samples in heritability estimates. Darkgray is for one kinship estimated by Kindred, and light gray is for kinship estimated by sample correlation matrix. The downsampling proportions are marked below violin plots. The total number of samples is 3925.
